# Methylphenidate boosts choices of mental labor over leisure depending on baseline striatal dopamine

**DOI:** 10.1101/859637

**Authors:** Lieke Hofmans, Danae Papadopetraki, Ruben van den Bosch, Jessica I. Määttä, Monja I. Froböse, Bram. B. Zandbelt, Andrew Westbrook, Robbert-Jan Verkes, Roshan Cools

## Abstract

The cognitive enhancing effects of methylphenidate are well established, but the mechanisms remain unclear. We recently demonstrated that methylphenidate boosts cognitive motivation by enhancing the weight on the benefits of a cognitive task in a manner that depended on striatal dopamine. Here we considered the complementary hypothesis that methylphenidate might also act by changing the weight on the opportunity cost of a cognitive task. To this end, fifty healthy participants (25 women) completed a novel cognitive effort discounting task that was sensitive to opportunity cost, and required choices between task and leisure. They were tested on methylphenidate, sulpiride or placebo and also underwent an [^18^F]DOPA PET scan to quantify baseline dopamine synthesis capacity. Methylphenidate boosted choices of cognitive effort over leisure across the group, and this effect was greatest in participants with more striatal dopamine at baseline. The effects of sulpiride did not reach significance. This study strengthens the motivational account of methylphenidate’s effects on cognition and suggests that methylphenidate reduces the cost of mental labor by increasing striatal dopamine.

## INTRODUCTION

The brain catecholamines have long been implicated in a wide range of cognitive functions, including working memory and cognitive control [1–3]. Drugs altering catecholamine transmission are first-line treatment for disorders accompanied by deficits in working memory and cognitive control, such as attention deficit/ hyperactivity disorder (ADHD) [4, 5] and are commonly used for cognitive enhancement in healthy people [6–8]. Various studies have demonstrated that acute administration of psychostimulants, like the dopamine and noradrenaline transporter blocker methylphenidate, enhances working memory and cognitive control and decreases feelings of fatigue in healthy individuals [9–16].

Such cognitive effects of catecholaminergic drugs have been most commonly attributed to a modulation of the ability to implement cognitive control, often associated with the prefrontal cortex [1]. However, recent progress suggests that cognitive control might also be altered by changing motivation, that is the willingness to engage with a cognitive task, rather than ability alone [17, 18]. Specifically, we have posited that the cognitive enhancing effects of drugs like methylphenidate, which act by blocking the dopamine and noradrenaline transporters, reflect changes in cost/benefit-based decision making about cognitive control, elicited by striatal dopamine [19, 20]. While prior evidence, for example from medication withdrawal studies in Parkinson’s disease, generally concurred with this hypothesis [18, 21–23] (but see [24]), there was, until recently, no direct evidence for a specific role for dopamine in the striatum. To definitively test this role for striatal dopamine in cognitive motivation, we set up two separate cognitive effort discounting experiments in the context of a large pharmacological PET study with 100 healthy volunteers. In this study we directly quantified striatal dopamine synthesis capacity with PET, while also measuring effects of methylphenidate and sulpiride. In both experiments, participants completed a working memory task prior to drug administration and a cognitive effort discounting task after drug administration, allowing us to isolate drug effects on motivation in a manner that was not confounded by drug effects on performance. In a separate session, participants underwent an [^18^F]DOPA PET scan to quantify baseline dopamine synthesis capacity. Uptake of the radiotracer [^18^F]DOPA indexes the degree to which dopamine is synthesized in (the terminals of) midbrain dopamine neurons, providing a relatively stable trait index of dopamine transmission that is less sensitive to state-dependent changes in dopamine levels [25] (but see [26]) than other dopamine PET tracers such as [^11^C]raclopride or [^18^F]fallypride, which reflect D2/3-receptor availability. To substantiate the hypothesis that the effects of the non-specific catecholamine enhancer methylphenidate (which increases both dopamine and noradrenaline in both striatum and cortex) reflect modulation of striatal dopamine, we compared the effects of methylphenidate with the effects of the selective D2-receptor antagonist sulpiride, which acts primarily on the striatum where D2-receptors are disproportionately abundant [27–29].

The two experiments in this large overarching pharmacological PET study were set up to test two complementary hypotheses about dopamine’s role in cognitive effort. The first experiment was inspired by neurocomputational modeling work of striatal dopamine (Opponent Actor Learning: OpAL model) [30], according to which striatal dopamine increases the weight on the benefit versus cost of options by shifting the balance of activity towards the direct Go pathway away from the indirect NoGo pathway of the basal ganglia. To test this hypothesis, half of the participants included in our study completed an experiment that we recently reported in Westbrook *et al*. [31], where participants chose between high-effort and low-effort options while we tracked their eye gaze. In line with the OpAL model [30], this experiment demonstrated that both methylphenidate and sulpiride boosted the selection of a high-versus low-effort task by increasing the weight on the benefits (monetary payoff) of the high-effort task. This drug effect was present only in participants with lower baseline levels of striatal dopamine, in line with the hypothesis that dopaminergic drug effects depend on variability in baseline levels of striatal dopamine [2, 22, 32].

The other half of the participants included in the large pharmacological PET study completed the experiment reported here. This experiment was motivated by a different hypothesis, derived from the recent opportunity cost theory of cognitive effort [33, 34], stating that performance of cognitive control tasks is costly, because it requires task focus and persistent task engagement, which interferes with performing potentially rewarding alternative tasks. Inspired by the proposal that the opportunity cost of physical effort, equal to the average reward rate of the environment, corresponds to levels of tonic dopamine [35, 36], the opportunity cost of cognitive effort was argued to also be carried by tonic dopamine [19, 33] (but see [37, 38]). To test this hypothesis, the current paradigm maximizes sensitivity to the opportunity cost of task engagement by allowing participants to choose between task engagement and leisure (allowing pursuit of unstructured/unspecified opportunities). By contrast, our previous experiment reported in Westbrook *et al*. [31] required choices between high- and low-effort options, thus controlling for opportunity cost.

For the present experiment, we considered two alternative hypotheses. First, we reasoned that prolonging the action of dopamine in the synapse via methylphenidate might potentiate task disengagement by amplifying a putatively dopamine-mediated signal of the opportunity costs of cognitive task engagement [19]. By contrast, we also considered the hypothesis that, in line with the OpAL model, methylphenidate might potentiate task engagement by shifting the balance more towards the benefits and away from the costs of cognitive work [30]. Given prior evidence for large individual variability in dopaminergic drug effects, we anticipated that this effect would depend on baseline striatal dopamine synthesis capacity. These predictions were preregistered on https://osf.io/g2z6p/.

## MATERIALS AND METHODS

### Participants

Fifty right-handed, neurologically and psychiatrically healthy volunteers (18-43 years at start of study, mean(SD) = 24(5.7); 25 women) were recruited as part of a larger study (detailed study overview in Supplementary information). Participants provided written informed consent and were paid €309 upon completion of the study. The study was approved by the local ethics committee (CMO region Arnhem-Nijmegen, The Netherlands: protocol NL57538.091.16; trial register NTR6140, https://www.trialregister.nl/trial/5959). One participant dropped out during the second day due to nausea, another after four study days due to anxiety, and PET data of two other participants were incomplete (one due to scanner software problems and another due to discomfort during scanning). We analyzed data of the resulting 46 participants.

### General study overview and pharmacological manipulation

The study consisted of five sessions. The first day served as an intake session. On the following three pharmacological sessions, participants first completed a working memory delayed response task (24 minutes), then they were administered an oral dose of either 20mg of methylphenidate, or 400mg of sulpiride, or placebo. Participants completed a cognitive effort-discounting choice procedure (22 minutes) 50 minutes after methylphenidate administration and 140 minutes after sulpiride administration. Methylphenidate plasma concentrations were expected to peak after 2 hours [39] and sulpiride plasma concentrations after 3 hours [40]. Drug timings were optimized for peak effects during an fMRI paradigm not reported here; near-peak effects were expected during the choice procedure. On the fifth day, participants underwent an [^18^F]DOPA PET scan to quantify their baseline dopamine synthesis capacity. See Supplementary information for complete task battery and timings.

### Behavioral paradigm

#### Color wheel working memory task

The color wheel task (Figure 1A) is a delayed response working memory task assessing two distinct component processes of cognitive control: distractor resistance and flexible updating [41, 42]. A more detailed description of the paradigm and a discussion on flexibility versus stability are reported in the Supplementary information. The primary research question of this study concerned drug effects on the motivation irrespective of the type of cognitive control process. On each trial, participants had 0.5s to memorize the colors and locations of one to four squares (set-size 1-4), followed by a 2s fixation cross. Then, a new set of colors appeared on screen for 0.5s, accompanied by either the letter ‘I’ (for ‘ignore’) or the letter ‘U’ (for ‘update’). In the ignore task-type, participants had to ignore the new colors and keep the previous set in memory. In the update task-type, they had to update their memory with the new set of colors. This was again followed by a fixation cross, which, depending on the task-type, lasted either 2s or 4.5s, ensuring equal delay times between the relevant stimuli (first set for the ignore type and second set for the update type) and the subsequent probe. During this probe phase, participants had 4s to indicate the color of the target square by clicking on the corresponding color on a color wheel. Participants completed 128 trials divided over 2 blocks, with an equal division across set-sizes and task-types.

**Figure 1.**
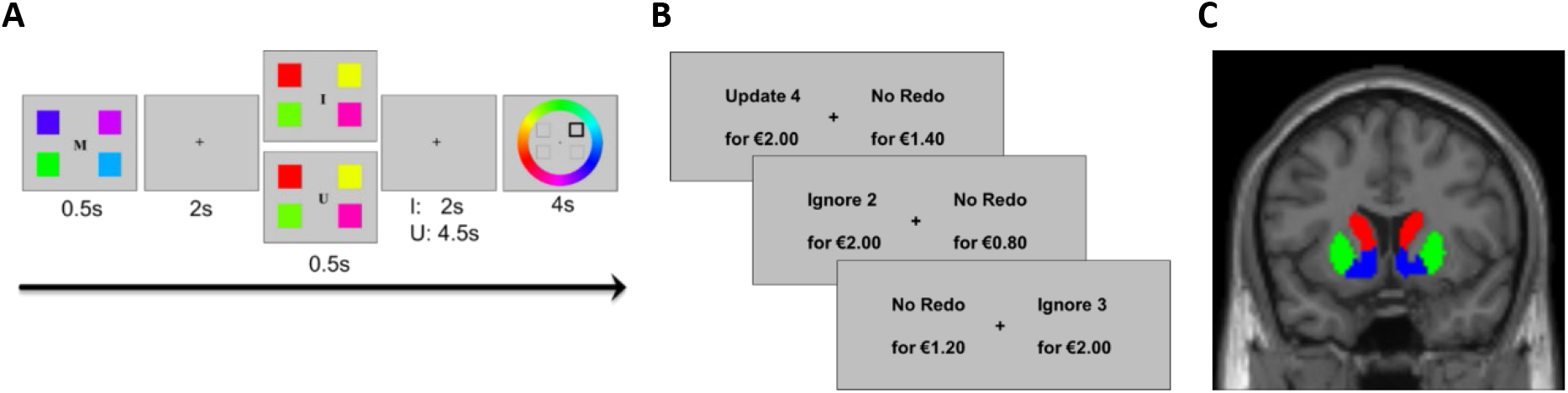
**A** – Schematic of the color wheel working memory task. I = ‘ignore’: participants have to ignore the new squares while still remembering the previous set of squares. U = ‘update’: participants have to remember the new set of squares and forget the previous set. **B** – Example trial sequence of the cognitive effort discounting choice task. **C** – Coronal view of our regions of interest including the nucleus accumbens (blue), putamen (green) and caudate nucleus (red).

#### Choice task

To quantify participants’ cognitive motivation, participants completed a choice task (Figure 1B) where they successively chose between repeating the color wheel task (redo option) for more money or a no-redo (rest) option for less money, in which participants would be free to do what they wanted for an equal length of time while staying in the testing room. Participants were informed that one of their choices would be randomly selected. Due to time constraints, and known to the participant, both the monetary bonus and the redo of the color wheel task were hypothetical. A strong effect of set-size on proportion of redo choices validated the task manipulation, evidencing strong monotonic cognitive load-based discounting (see Results). The hypothetical compensation for the redo option was fixed at €2.00. The compensation for the no-redo option ranged from €0.10 to €2.20. The redo option was further specified by task-type and set-size, so that participants were instructed that the majority of trials in the redo block would consist of trials of the chosen task-type and set-size. The remainder of the trials would be randomly divided among all task-type and set-size combinations. To account for the stochastic nature of decision-making [43], we opted not to use a titration procedure for arriving at the subjective value [44], since titration adjusts the offer for the no-redo option based on previous, noisy choices. Instead, we randomly sampled choices across the full value-range in 3 equivalent blocks of 96 trials each, equally divided across set-size, task-type and monetary offer for the no-redo option.

### PET acquisition and preprocessing

PET scans were acquired on a Siemens PET/CT-scanner at the Department of Nuclear Medicine of the Radboudumc, using an [^18^F]DOPA radiotracer, produced by the Radboud Translational Medicine department. Participants received 150mg carbidopa and 400mg entacapone 50 minutes before scanning, to minimize peripheral metabolism of [^18^F]DOPA by decarboxylase and COMT, respectively, thereby increasing signal to noise ratio in the brain. After a bolus injection of [^18^F]DOPA (185MBq; approximately 5mCi) into the antecubital vein, the procedure started with a low dose CT-scan (approximately 0.75mCi) to use for attenuation correction of the PET images after which a dynamic PET scan was collected over 89 minutes and divided into 24 frames (4×1, 3×2, 3×3, 14×5 min). PET data (4×4×3mm voxel size; 5mm slice thickness; 200×200×75 matrix) were reconstructed with weighted attenuation correction and time-of-flight recovery, scatter-corrected, and smoothed with a 3mm full-width-at-half-maximum (FWHM) kernel. For registration purposes we acquired a T1-weighted anatomical MRI scan on the first testing day, using an MP-RAGE sequence (repetition time = 2300ms, echo time = 3.03ms, 192 sagittal slices, field of view = 256mm, voxel size 1mm isometric) on a Siemens 3T MR-scanner with a 64-channel coil. After reconstruction, PET data were preprocessed using SPM12 (http://www.fil.ion.ucl.ac.uk/spm/). All frames were realigned for motion correction and coregistered to the anatomical MRI-scan, using the mean PET image of the first 11 frames (using the mean image of only the first 11 frames improves coregistration, because these images have a greater range in image contrast in regions outside the striatum). Dopamine synthesis capacity was computed per voxel as [^18^F]DOPA influx constant per minute (K_i_) relative to the cerebellar grey matter reference region using Gjedde-Patlak graphical analysis on the PET frames from the 24th to 89th minute [45]. We then extracted average K_i_ values from three regions of interest (ROIs) – nucleus accumbens, putamen and caudate nucleus – defined using masks based on an independent functional connectivity-analysis of the striatum [46] and exactly the same as reported in Westbrook *et al*. [31] (Figure 1C).

### Data analysis

Performance measures on the color wheel task included median absolute degrees of deviance of the response from the correct color (deviance) and median response time (RT) for each participant. Participants’ preferences on the choice task were calculated as the proportion of trials on which participants chose the redo option over the no-redo option (proportion redo). Outliers were a priori defined as those who deviated more than three standard deviations from the global mean, which did not result in any exclusions. Data were analysed in JASP (version 0.11.1) using separate repeated-measures ANOVAs for each ROI – nucleus accumbens, putamen and caudate nucleus, including drug (placebo, methylphenidate, or sulpiride), task-type (ignore or update) and set-size (ranging from 1 to 4) as within-subjects variables and baseline dopamine synthesis capacity (measured as the mean-centered average [^18^F]DOPA uptake, K_i_) as covariate. Greenhouse–Geisser corrections were applied when the sphericity assumption was violated. A *p*-value smaller than 0.017 (Bonferroni-corrected for the 3 ROIs) was considered significant.

## RESULTS

### Working memory performance

Before drug intake, participants performed the working memory task. Across sessions and in line with earlier work [42], participants performed poorer when working memory load increased, as indicated by higher deviance (*F*_(1.55,68.30)_ = 37.1, *p* < 0.001) and longer RTs (*F*_(2.35,103.44)_ = 143.6, *p* < 0.001). While participants deviated from the target color less on update trials (*F*_(1,44)_ = 44.7, *p* < 0.001), their RTs were longer compared with ignore trials (*F*_(1,44)_ = 31.0, *p* < 0.001). Both deviance and RT show a significant interaction (deviance: *F*_(1.67,73.60)_ = 14.7, *p* < 0.001; RT: *F*_(2.71,119.24)_ = 4.4, *p* = 0.007; Figure 2A-B).

**Figure 2.**
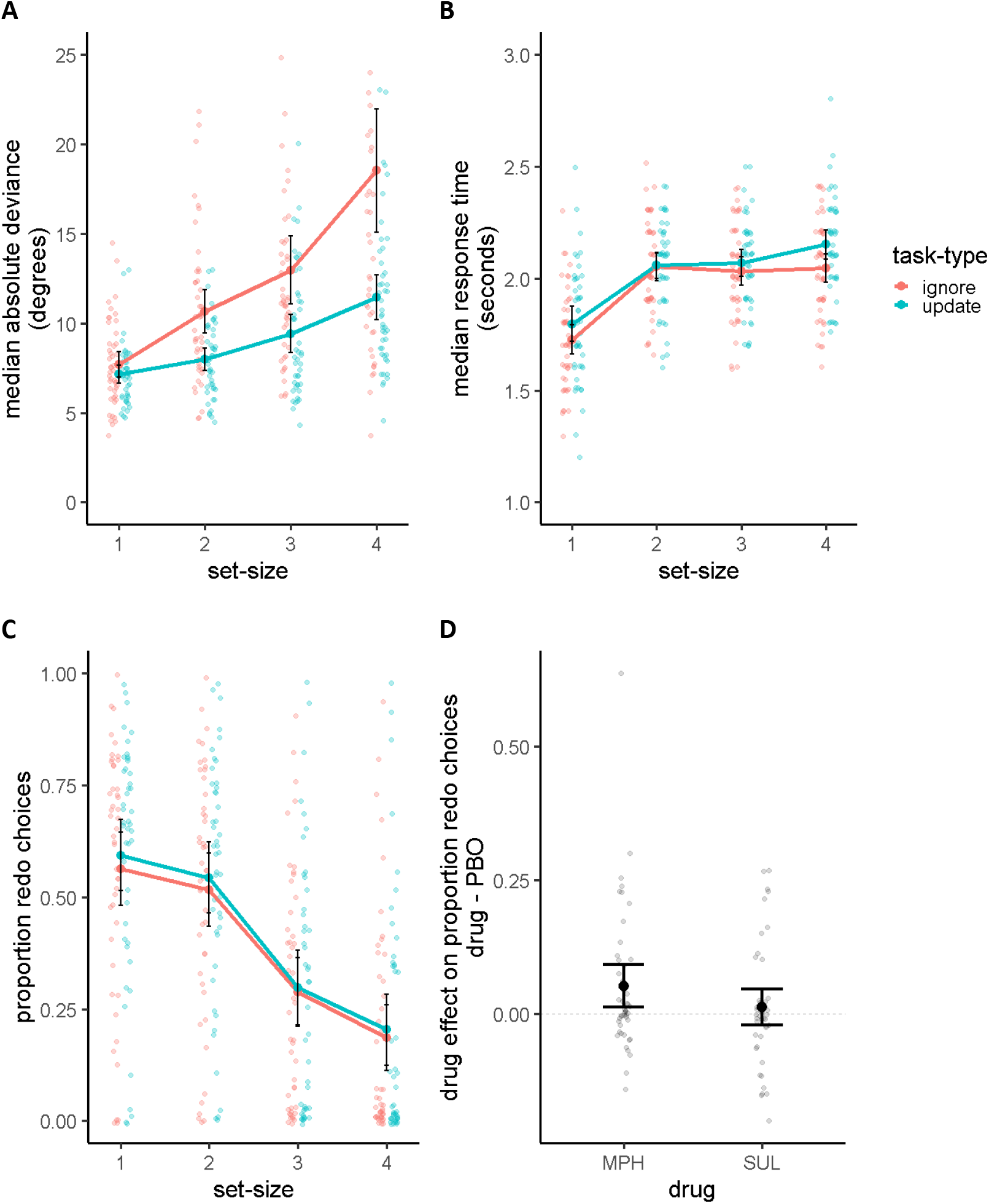
**A** – Median absolute deviance, **B** – median response times and **C** – the proportion of trials on which participants chose the redo option across drug sessions plotted as a function of set-size, separately for each task-type. **D –** Drug effect on the proportion of trials on which participants chose the redo option (methylphenidate or sulpiride minus placebo). The methylphenidate-induced effect on proportion redo choices is still significant without the participant showing the greatest effect: *F*_(2,86)_ = 4.0, *p* = 0.022. Error bars represent 95% confidence interval around the mean. MPH: methylphenidate; SUL: sulpiride; PBO: placebo.

There was no main effect of dopamine synthesis capacity on either deviance (caudate nucleus: *F*_(1,44)_ = 0.5, *p* = 0.471; putamen: *F*_(1,44)_ = 0.1, *p* = 0.795; nucleus accumbens: *F*_(1,44)_ = 0.2, *p* = 0.634) or RT (caudate nucleus: *F*_(1,44)_ = 0.04, *p* = 0.838; putamen: *F*_(1,44)_ = 0.3, *p* = 0.571; nucleus accumbens: *F*_(1,44)_ = 0.2, *p* = 0.652), nor did dopamine synthesis capacity interact with any of the other variables.

### Methylphenidate increased cognitive motivation

At baseline, participants exhibited a preference for not repeating any task, as evidenced by the overall proportion redo being significantly smaller than 0.5 on the placebo session (proportion = 0.38, SD = 0.24; *t*_(45)_ = −3.5, *p* = 0.001). As hypothesized, we found a significant effect of drug on proportion redo (main effect of drug with 3 conditions: *F*_(2,88)_ = 5.1, *p* = 0.008). This was driven by higher proportion redo under methylphenidate versus placebo (*F*_(1,44)_ = 8.0, *p* = 0.007; Figure 2D). There was no difference between sulpiride and placebo (*F*_(1,44)_ = 0.6, *p* = 0.444; Figure 2D). Numerically, proportion redo was higher under methylphenidate than sulpiride, but this difference did not survive correction for multiple comparisons (*F*_(1,44)_ = 5.7 *p* = 0.021). Proportion redo decreased with set-size (*F*_(1.37,60.17)_ = 79.3, *p* < 0.001; Figure 2C). There was no effect of task-type (*F*_(1,44)_ = 1.3, *p* = 0.268) and no interaction between task-type and set-size (*F*_(2.18,95.81)_ = 1.4, *p* = 0.247), nor did drug interact with task-type (*F*_(1.60,70.59)_ = 0.7, *p* = 0.479) or set-size (*F*_(3.00,132.19)_ = 0.8, *p* = 0.496).

### High-dopamine participants exhibited greater methylphenidate-related increases in cognitive motivation

The effect of methylphenidate on proportion redo depended on baseline levels of dopamine synthesis capacity. This was supported by a significant interaction between drug (methylphenidate, sulpiride, placebo) and dopamine synthesis capacity in the nucleus accumbens (*F*_(2,88)_ = 5.0, *p* = 0.009; Figure 3B-C; Table 1). Participants with higher dopamine synthesis capacity in the nucleus accumbens exhibited greater methylphenidate-induced increases in proportion redo choices than participants with lower dopamine synthesis capacity (*F*_(1,44)_ = 8.3, *p* = 0.006). The drug by dopamine synthesis capacity interaction for sulpiride versus placebo was not significant, while there was a trend towards a drug by dopamine synthesis capacity interaction for methylphenidate versus sulpiride (*F*_(1,44)_ = 4.8, *p* = 0.034). Although sub-threshold, interactions in the same direction were found between drug and dopamine synthesis capacity in the putamen and caudate nucleus (Figure 3E-F,H-I; Table 1). There was also a sub-threshold negative association between baseline dopamine synthesis capacity and proportion redo choices under placebo in the nucleus accumbens, but not the other ROIs (Figure 3A,D,G; Table 1). Importantly, supplementary analyses demonstrated that the effect of methylphenidate on proportion redo does not reflect changes in choice randomness, or effects on task performance (completed prior to drug administration), or effects on mood and medical symptoms, and are reproduced when analyzing ‘indifferent points’ (Supplementary information).

**Figure 3.**
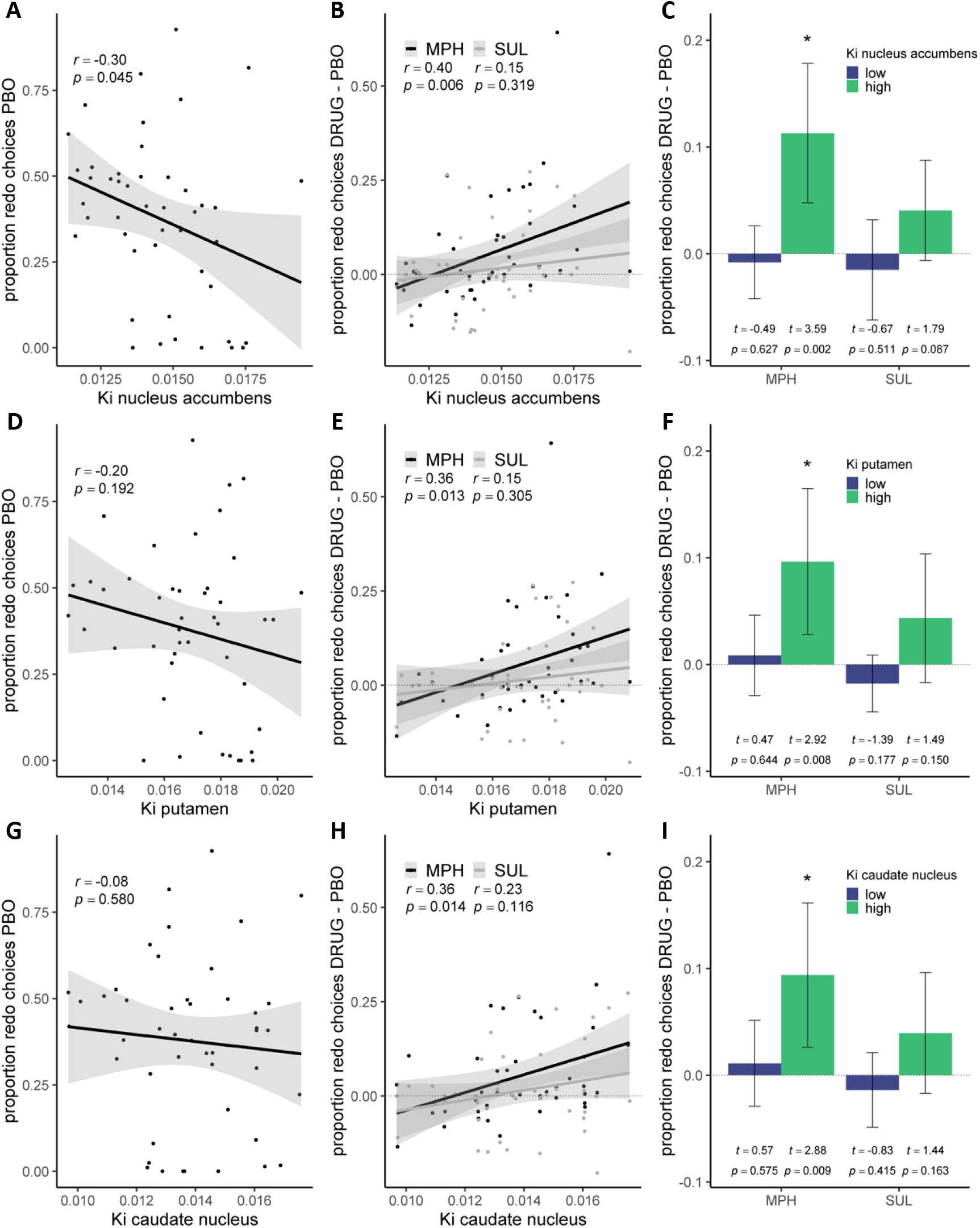
Proportion redo choices as a function of dopamine synthesis capacity in the nucleus accumbens (upper panels), putamen (middle panels) and caudate nucleus (bottom panels). *p*-values < 0.017 were considered significant. **A.** Correlation between dopamine synthesis capacity and proportion redo choices under placebo. **B.** Correlation between dopamine synthesis capacity and drug-induced changes in proportion redo choices. Correlation coefficients and *p*-values without the participant showing the greatest methylphenidate-induced effect on proportion redo choices: *r_nucleus accumbens_* = 0.36, *p* = 0.014; *r_putamen_* = 0.41, *p* = 0.005; *r_caudate nucleus_* = 0.29, *p* = 0.056. **C.** Median split on dopamine synthesis capacity for visualization purposes. Shaded areas and error bars represent 95% confidence interval around the mean. PBO = placebo; MPH = methylphenidate; SUL = sulpiride; K_i_ = [^18^F]DOPA uptake.

**Table 1.**
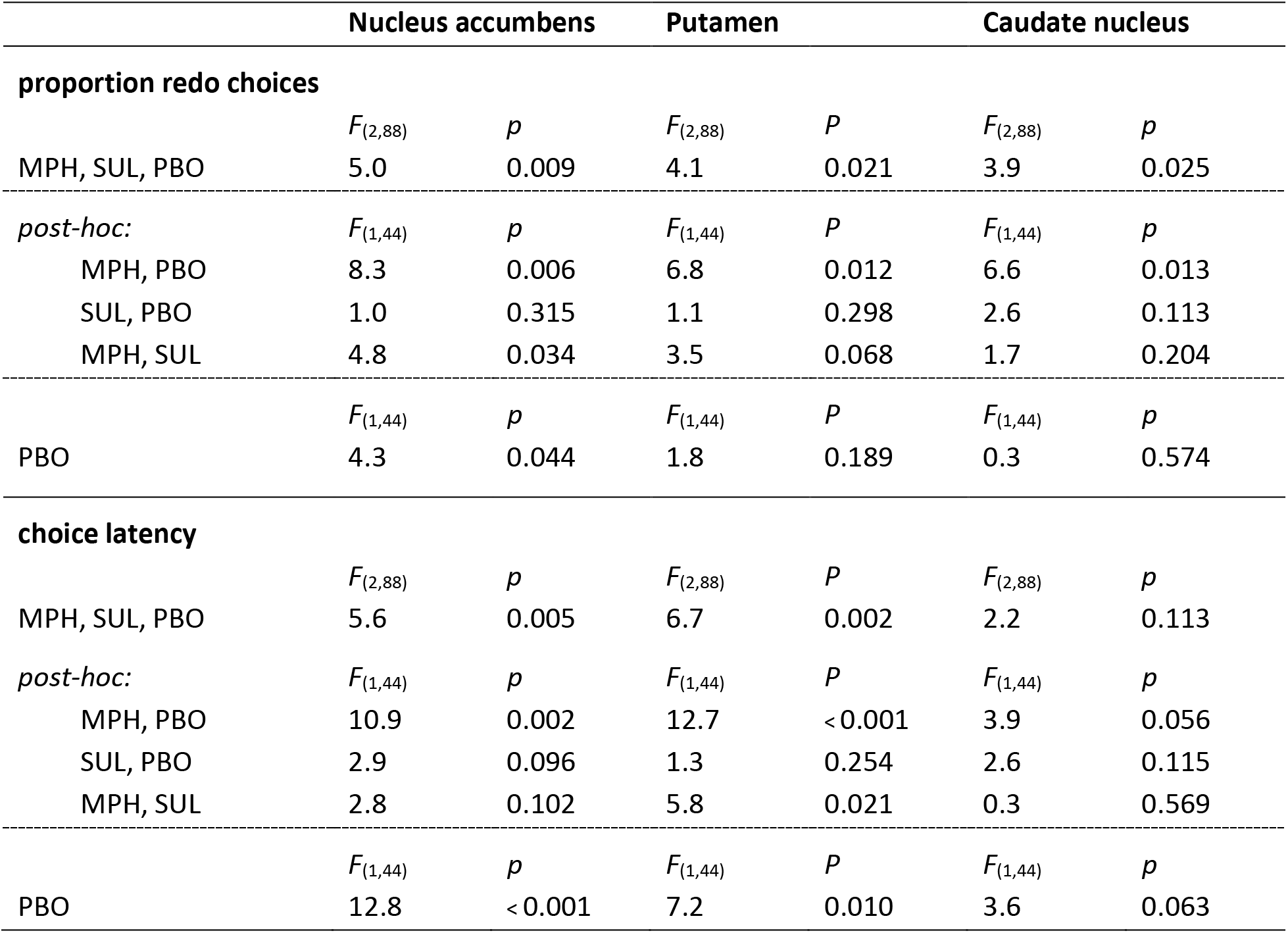
Repeated-measures ANOVAs on proportion redo choices and choice latency. Separate analysis for each ROI – nucleus accumbens, putamen and caudate nucleus, including drug, set-size and task-type as within-subjects variables and baseline dopamine synthesis capacity (measured as the mean-centered average [^18^F]DOPA uptake, K_i_) as covariate. Statistics for the interaction between baseline dopamine synthesis and drug are shown, as well as the main effect of dopamine synthesis capacity on the placebo session. MPH = methylphenidate; SUL = sulpiride; PBO = placebo. *p*-values below a Bonferroni-corrected alpha-value of 0.017 were considered significant.

### Drug manipulation does not interact with the benefit of engaging in a cognitive task

Primary analyses on proportion redo choices revealed no significant interactions between drug and the cognitive cost of the task – the set-size. We also explored whether drug effects interacted with the benefit of the task – the monetary payoff for the redo option relative to the no-redo option. Note that the payoff of the redo options was constant throughout the task. To that end, we added, in an additional analysis, the monetary payoff for the no-redo option to our rmANOVA. Because each monetary value was only repeated three times per drug session, task-type and set-size, we divided these values into tertiles so that proportion redo was calculated based on 12 trials. As expected, the monetary payoff had a strong negative main effect on proportion redo (*F*_(2,88)_ = 114.9, *p* < 0.001), such that the higher the payoff for the no-redo option, the less often people chose the redo option. Although numerically there was a greater methylphenidate-related increase in proportion redo when the payoff for the no-redo option was lower (i.e. when the benefit for the task was higher), payoff did not significantly interact with drug (*F*_(4,176)_ = 2.4, *p* = 0.062) or with the interaction of drug with dopamine synthesis capacity (nucleus accumbens: *F*_(4,176)_ = 0.7, *p* = 0.561; putamen: *F*_(4,176)_ = 0.8, *p* = 0.513; caudate nucleus: *F*_(4,176)_ = 0.9, *p* = 0.456). Thus, while the possibility of unstructured free time comprised unmeasured opportunity costs of engaging in the task, drug manipulation did not reliably affect the sensitivity to the relative, explicit costs or benefits of the redo option.

### High-dopamine participants exhibited greater methylphenidate-related slowing of choice latency

Exploratory analyses of choice latency revealed no main effect of drug (*F*_(2,88)_ = 0.2, *p* = 0.797). There was a significant interaction between effect of drug on choice latency and dopamine synthesis capacity in the nucleus accumbens (*F*_(2,88)_ = 5.6, *p* = 0.005) and putamen (*F*_(2,88)_ = 6.7, *p* = 0.002; Table 1), which was driven by a difference between methylphenidate and placebo (nucleus accumbens: *F*_(1,44)_ = 10.9, *p* = 0.002; putamen: *F*_(1,44)_ = 12.7, *p* < 0.001; Table 1; Figure 4B-C,E-F,H-I). Methylphenidate slowed people with higher baseline dopamine synthesis capacity and invigorated people with lower baseline dopamine synthesis capacity. No significant interaction between dopamine synthesis capacity and the effect of sulpiride, or between dopamine synthesis capacity and the difference between methylphenidate and sulpiride was observed (Table 1). A negative association between baseline dopamine synthesis capacity and choice latency under placebo was present in the nucleus accumbens (*F*_(1,44)_ = 12.8, *p* < 0.001) and putamen (*F*_(1,44)_ = 7.2, *p* = 0.010), and trending in the caudate nucleus (*F*_(1,44)_ = 3.6, *p* = 0.063), indicating that higher baseline dopamine synthesis capacity was associated with faster responding (Figure 4A,D,G; Table 1).

**Figure 4.**
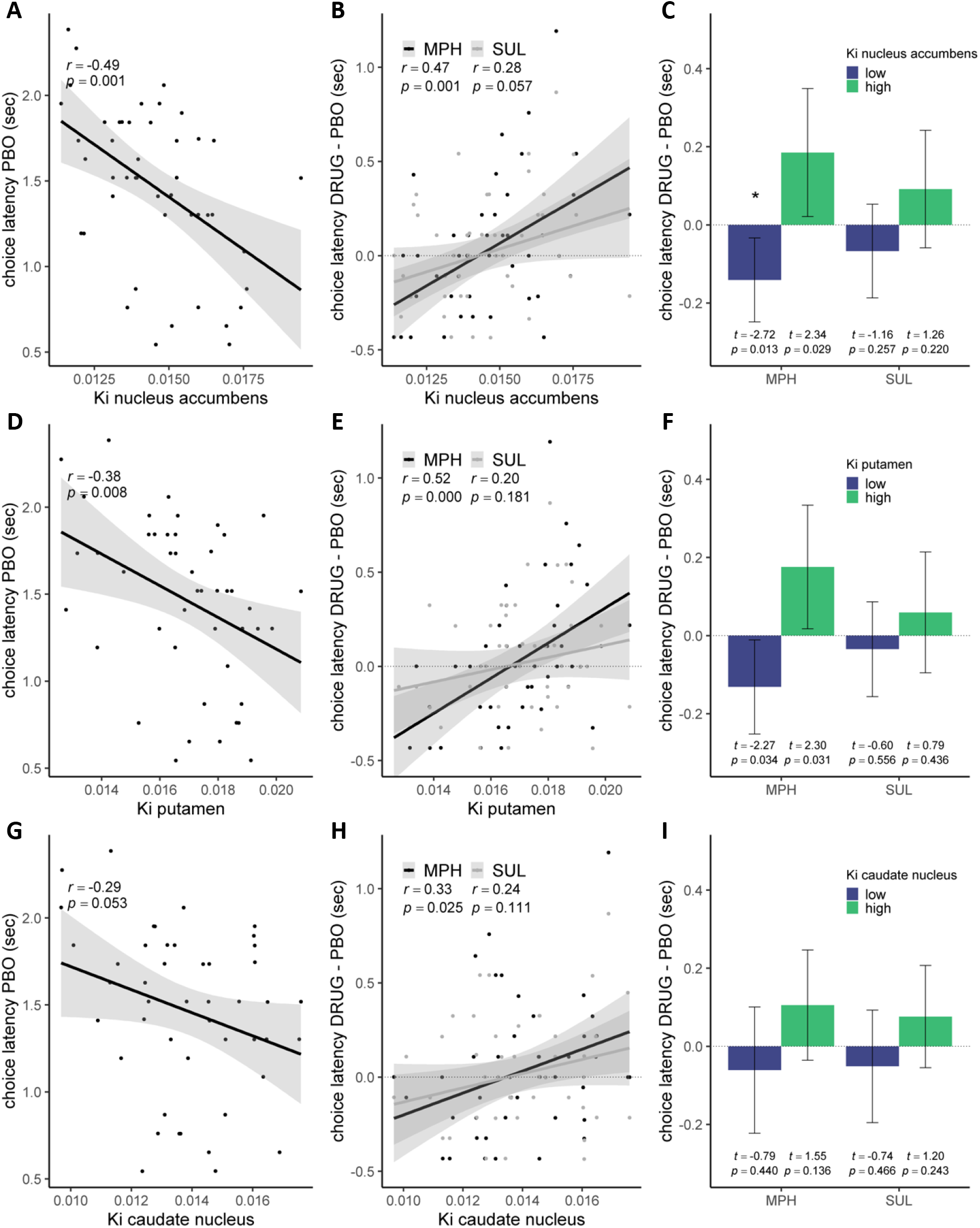
Choice latency as a function of dopamine synthesis capacity in the nucleus accumbens (upper panels), putamen (middle panels) and caudate nucleus (bottom panels). *p*-values < 0.017 were considered significant. **A.** Correlation between dopamine synthesis capacity and choice latency under placebo. **B.** Correlation between dopamine synthesis capacity and drug-induced changes in choice latency. **C.** Median split on dopamine synthesis capacity for visualization purposes. Shaded areas and error bars represent 95% confidence interval around the mean. PBO = placebo; MPH = methylphenidate; SUL = sulpiride; K_i_ = [^18^F]DOPA uptake.

### Positive correlation between drug-induced effects on cognitive motivation and choice latency

Individuals who showed greater methylphenidate-related increases in proportion redo also showed greater methylphenidate-related slowing (Pearson’s *r* = 0.67, *p* < 0.001). A similar positive correlation was present between the effect of sulpiride versus placebo on choice latency and the drug effect on proportion redo (*r* = 0.50, *p* < 0.001).

All region-of-interest based results were corroborated by voxel-wise K_i_ analyses and were present after controlling for session order (Supplementary information).

## DISCUSSION

The present study demonstrates that methylphenidate boosts motivation for cognitive task performance over leisure. This effect was present across the group as a whole but was particularly strong in people with high ventral striatal dopamine synthesis capacity. This finding is consistent with the OpAL model [30], stating that methylphenidate reduces the weight on the cost of task engagement. Together with the findings reported in Westbrook *et al*. [31], these data strengthen the link between striatal dopamine and cognitive motivation [18, 47] and the hypothesis that the cognitive enhancing effect of methylphenidate reflects an increase in motivation. The present study design provides a particularly good test of drug-induced changes in participants’ cognitive motivation, rather than capacity, because methylphenidate was administered after the task-performance phase, but before the discounting phase. Moreover, the data firmly establish the pervasive baseline-dependency hypothesis of individual variability in the efficacy of the most commonly used catecholaminergic drug, methylphenidate.

The present paradigm was more sensitive to the motivational boosting effect of methylphenidate, which was observed across the group as a whole, than the paradigm in Westbrook *et al*. [31], where the effect was detected only in low-dopamine participants. This likely reflects the greater sensitivity of the current paradigm, at baseline, to task avoidance, as evidenced by a strong preference for the rest option. We argue that this increased sensitivity to task avoidance of the present paradigm reflects the increased opportunity cost of task engagement: By choosing the task option, they also chose to forego an opportunity to rest and play with their smartphone and/or laptop. This sensitivity to the opportunity cost at baseline, which tended to be greater in high-dopamine participants, might have rendered greater dynamic range for methylphenidate-related decreases in the weight on the cost. Conversely, Westbrook *et al*. required choices between a high effort option for more money and a low effort option for less money. This set-up controlled for opportunity costs, and generated a default preference for the high-reward high-effort task. This higher preference for the effortful option at baseline, particularly in high-dopamine participants, might have reduced the range for further increases in the weight on the benefits in those participants. In short, the two paradigms likely differ in their sensitivity to increases in the benefits versus decreases in the costs by methylphenidate. This is supported by the finding that the effect of methylphenidate in the previous experiment, but not the current experiment, interacted with monetary payoff. Critically, the differential sensitivity of the two paradigms to changes in the benefits versus (opportunity and/or effort) costs of cognitive effort might also underlie the observation that methylphenidate effects are greater in high-dopamine participants in the present experiment but, conversely, in low-dopamine participants in Westbrook *et al*. Future studies might address the question whether the different types of effort costs and benefits implicate dopamine in distinct subregions of the striatum. This hypothesis is raised cautiously by the finding that the effect of methylphenidate on effort selection in the present study depends most strongly on dopamine synthesis capacity in the nucleus accumbens, whereas the effect in Westbrook *et al*. depends most strongly on dopamine in the caudate nucleus.

The finding of a strong negative association between choice latency and dopamine synthesis capacity under placebo concurs with the well-established link between dopamine and response vigor [48–50]. However, while methylphenidate sped up choices of participants with low baseline dopamine synthesis capacity, it slowed choices of participants with higher dopamine synthesis capacity. Intriguingly, these baseline-dependent effects of methylphenidate on choice latency correlated with the effects of methylphenidate on cognitive effort choice. Therefore, one explanation of the seemingly conflicting effect of methylphenidate on choice latency is that the strength of the default preference for no-redo was strongest for people with high dopamine synthesis capacity. Because these participants showed the largest shift away from a default preference on methylphenidate, they might have experienced greater choice conflict, accounting for their slowing.

A limitation of the current large pharmacological PET study is that it does not allow us to directly address the neural locus of methylphenidate’s effect. The finding that the effects of methylphenidate were associated with striatal dopamine synthesis capacity suggests that methylphenidate acted on the striatum to modulate cognitive motivation. However, given that [^18^F]DOPA uptake signal is too low in the prefrontal cortex, we cannot exclude the possibility that the variation in nigrostriatal dopamine synthesis capacity is paralleled by variation in prefrontal dopamine levels. An additional prefrontal locus of effect is also consistent with the absence of significant effects of sulpiride, which acts selectively on dopamine D2-receptors that are particularly abundant in the striatum. In future studies pharmacology and PET should be combined with functional magnetic resonance imaging to isolate the neural locus of the baseline-dependent effects of methylphenidate on cognitive motivation.

The finding that the effects of methylphenidate, which blocks both dopamine and noradrenaline transporters [51, 52], were not accompanied by significant effects of the selective D2-receptor antagonist sulpiride is surprising. First, previous research has established that the present dose of sulpiride is effective at approximately 2 hours after intake, indexed in terms of both sulpiride plasma concentrations [40, 53] and behavioral effects on reversal learning [54]. Second, the exact same dose of sulpiride did have a significant effect in Westbrook *et al*. [31], where the exact same study protocol was applied. This might lead some to ask whether the current effects reflect a modulation of noradrenaline instead of dopamine [55–59]. However, given the lack of sulpiride-related changes in physiological or subjective report measures (Supplemental information), we cannot exclude the possibility that the drug manipulation did not alter behavior on the current task due to, for example, mixing of pre- and postsynaptic effects, insufficient dosing and/or suboptimal timing of testing. Moreover, given that methylphenidate’s effects were predicted by [^18^F]DOPA uptake in the striatum (which does not contain any noradrenaline receptors), together with the presence of trend effects of sulpiride going in the same direction as those of methylphenidate, we argue that the effects reflect modulation of dopamine rather than noradrenaline. Indeed this conclusion also concurs with prior evidence that methylphenidate’s enhancing effects correspond with changes in midbrain dopamine release [60].

In conclusion, this study suggests that methylphenidate reduces the cost of mental labor by increasing striatal dopamine, thus strengthening the motivational account of methylphenidate’s effects on cognition.

## ACKNOWLEDGMENTS

We thank Britt Lambregts, Margot van Cauwenberge, Dirk Geurts, Peter Mulder and Monique Timmer for assistance during data collection.

## AUTHOR CONTRIBUTIONS

Conceptualization: R.C., methodology: R.C., D.P., M.I.F., B.B.Z., L.H., software: D.P., L.H., R.v.d.B., formal analysis: L.H., R.v.d.B., R.C., investigation: L.H., D.P., R.v.d.B., J.I.M., R-J.V., data curation: L.H., D.P., R.v.d.B., J.I.M., writing – original draft preparation: L.H., R.C., writing – review and editing: L.H., D.P., R.v.d.B., J.I.M, M.I.F., B.B.Z., A.W., R-J.V., R.C., visualization: L.H., B.B.Z., supervision: R.C., project administration: J.I.M., funding acquisition: R.C.

