## Supplemental information for "Methylphenidate boosts choices of mental labor over leisure depending on baseline striatal dopamine"

### SUPPLEMENTAL METHODS

#### Task battery

The current experiment was part of a larger study (*N* = 100). The first twenty-five women and twenty-five men completed the color wheel task. A description of the task battery, timings and randomization of the drug sessions can be found in Tables S1-3.

The study consisted of five sessions, with an interval of at least one week between each session. The first study day served as an intake on which participants were screened for inclusion criteria (see below), underwent an anatomical MR scan, and completed several baseline measures (Table S4). On the following three pharmacological sessions participants first completed a working memory delayed response task, then they received an oral administration of either 20 mg of methylphenidate, or 400 mg of sulpiride, or a placebo. Methylphenidate is a catecholamine-transporter blocker, which reduces the reuptake of catecholamines, including dopamine and norepinephrine, thereby increasing their availability in the synapse. Sulpiride is a selective D2 receptor antagonist that acts specifically on the dopamine system. Timings of drug administration were optimized for peak drug effects during the fMRI paradigm (not reported here; Table S2): methylphenidate plasma concentrations peak after 2 hours [1] and sulpiride plasma concentrations peak after 3 hours [2]. Near-peak effects were expected during the task of interest. After drug administration participants completed a cognitive effort-discounting choice procedure. Blood pressure, heart rate, and ear temperature were measured three times during the day for monitoring and safety reasons. At the same time-points, medical symptoms and mood measures were assessed three times on each session: at the start of each session, 20 minutes before the discounting task and at the end of the session (Table S5). On the fifth day, participants underwent an [$^{18}$F]DOPA PET scan to quantify their baseline dopamine synthesis capacity and completed several baseline measures (Table S4).

**Table S1. Task battery**

| DAY 1: INTAKE | DAY 2-4: PHARMACOLOGICAL-fMRI | HOME: QUESTIONNAIRES | DAY 5: PET SCAN |
|---|---|---|---|
| **HEALTH SCREENING** | **COGNITIVE ASSESSMENT** | **PERSONALITY ASSESSMENT** | **COGNITIVE ASSESSMENT** |
| **Psychiatric assessment** | **Executive functions** | Impulsivity (BIS-11A) | **Executive functions** |
| M.I.N.I Plus 5.0.0 | Color wheel working memory task[1] | Behavioral inhibition/activation | Digit span |
| **Physiological measures** | N-back working memory task[1] | Need for Cognition | **Intellectual functioning** |
| Electrocardiogram (ECG) | **Cognitive motivation** | Depression inventory (BDI) | Fluid intelligence (matrix reasoning) |
| Heart rate / blood pressure | Cognitive effort discounting task | Creativity scale (K-DOCS) | **Motivation/learning** |
| Body temperature | **Reward processing** | | Pavlovian to instrumental transfer task |
| Spontaneous eye blink rate | Reinforcement learning and working memory (RLWM) | | |
| **MRI** | **Creativity** | | **DOPAMINE SYNTHESIS CAPACITY** |
| Anatomical T1 scan | Alternative uses task | | **Positron emission tomography (PET)** |
| | Remote association task | | Carbidopa and entacapone intake |
| **COGNITIVE ASSESSMENT** | Alternative names task | | [$^{18}$F]DOPA bolus injection |
| **Executive functions** | | | |
| Digit span | **fMRI** | | |
| Listening span | **Reward processing and motivation** | | |
| **Intellectual functioning** | Reversal learning task (RL) | | |
| Crystalized IQ (NLV) | Monetary Incentive Delay task (MID) | | |
| | **SOMATIC MEASURES** | | |
| | Heart rate / blood pressure | | |
| | Body temperature | | |
| | **MOOD ASSESSMENT** | | |
| | Positive and negative affect scales (PANAS) | | |
| | Visual analogue scale (VAS) | | |

[1] First fifty participants completed the color wheel task (data reported here) while the second fifty completed the N-back task [3]. M.I.N.I.: Dutch M.I.N.I. International Neuropsychiatric Interview 5.0; NLV: Dutch reading test; BIS-11A: Barratt Impulsiveness Scale; BDI: Beck Depression Inventory II; K-DOCS: Kaufman Domains of Creativity Scale.

**Table S2. Pharmacological-fMRI session – timings**

| **Description** | **Sulpiride** | **Methylphenidate** |
|---|---|---|
| Screening | -70 | -160 |
| Somatic and mood measures | -65 | -155 |
| **BEH:** color wheel or N-back | -40 | -130 |
| **Capsule 1:** SUL/PBO | 0 | -90 |
| Rest | 1 | -89 |
| **Capsule 2:** MPH/PBO | 89 | 0 |
| **MRI:** screening | 105 | 15 |
| Somatic and mood measures | 120 | 30 |
| **BEH:** cognitive effort discounting task | 140 | 50 |
| **MRI:** installment | 170 | 80 |
| **MRI:** RL | 185 | 95 |
| **MRI:** MID | 215 | 125 |
| Lunch | 235 | 145 |
| **BEH:** RLWM | 250 | 160 |
| **BEH:** Creativity | 290 | 200 |
| Somatic and mood measures | 308 | 218 |
| End study day | 328 | 238 |

Timings are in minutes, where T = 0 is the time of drug intake. Participants either received sulpiride (SUL) followed by placebo (PBO), PBO followed by methylphenidate (MPH) or PBO twice in a within-subjects, cross-over double-blind design. BEH: behavioral testing; MRI: functional magnetic resonance imaging. RL: reversal learning task; MID: monetary incentive delay task; RLWM: reinforcement learning and working memory task.

**Table S3. Number of participants per drug order.**

| Day 2 | Day 3 | Day 4 | No. Participants |
|---|---|---|---|
| MPH | SUL | PBO | 10 |
| MPH | PBO | SUL | 10 |
| SUL | MPH | PBO | 6 |
| SUL | PBO | MPH | 7 |
| PBO | MPH | SUL | 6 |
| PBO | SUL | MPH | 7 |

PBO: placebo; MPH: methylphenidate; SUL: sulpiride

**Table S4. Demographic and background characteristics of participants.**

| | Characteristic | Measure | | | | |
|---|---|---|---|---|---|---|
| **Demographics** | Gender | Women / men (number) | 23/23 | | | |
| | | | **Min** | **Max** | **Mean** | **SD** |
| | Age | years | 18 | 43 | 23.8 | 5.9 |
| **Neuropsychological assessment** | Working memory capacity | Listening span: total span | 0 | 6.5 | 4.2 | 1.5 |
| | | Digit span | | | | |
| | | Forward | 4.5 | 12.5 | 8.2 | 1.9 |
| | | Backward | 3.5 | 11.5 | 7.3 | 2.1 |
| | Verbal intelligence | NLV[1] | 69.5 | 98.0 | 84.8 | 5.9 |
| | Fluid intelligence | WAIS-IV-NL | 9 | 22 | 16.8 | 3.1 |
| **Self-report questionnaires** | Trait impulsivity | BIS-11[2]: total score | 41.3 | 83.8 | 66.4 | 8.7 |
| | Need for Cognition | NCS | 41 | 81 | 61.2 | 9.0 |
| | Depressive symptoms | BDI | 0 | 13 | 3.7 | 3.4 |
| | Behavioral inhibition/activation | BIS/BAS: total score | 22 | 48 | 39.3 | 5.2 |
| | Creativity | K-DOCS: total score | 2.2 | 5 | 3.0 | 0.5 |

Minimum and maximum score, mean and standard deviation (SD) of demographic and background characteristics of participants included in the behavioral analyses. Neuropsychological assessment included the Dutch reading test (NLV; [4]), listening span task [5, 6], digit span test (forward and backward; [7]), and the matrix reasoning subtest of the WAIS-IV-NL [7]. Questionnaires included the Barratt Impulsiveness Scale (BIS-11A; [8]), Need for Cognition Scale (NCS; [9]), Beck Depression Inventory II (BDI-II; [10]), Behavioral Inhibition Scale/Behavioral Activation Scale (BIS/BAS; [11]) and the Kaufman Domains of Creativity Scale (K-DOCS; [12]). *N* = 46. [1]1 missing value. [2]The BIS-11A, an earlier version of the BIS-11 [13], was inadvertently sent to the participants. We converted the scores to BIS-11 [14] to be able to compare impulsivity scores with earlier studies.

Scores are comparable with earlier observations in healthy populations, e.g. Froböse et al., 2018 [15]: Listening span: mean = 4.8; Digit span forward: mean = 8.3; Digit span backward: mean = 7.2; BIS-11: mean = 61.8; NCS: mean = 63.3; BDI: mean = 3.6; BIS/BAS: mean total score = 39.7. Cacciaglia et al., 2018 [16]: WAIS-IV matrix reasoning: mean = 16.3. Pretz & Kaufman, 2015 [17]: K-DOCS total score: mean = ~3.2. Timmer et al., 2017 [18]: NLV-IQ mean = 97.8, equivalent to NLV mean = ~86.0.

**Table S5. Medical symptoms and mood measures**

| | PBO | | | MPH | | | SUL | | |
|---|---|---|---|---|---|---|---|---|---|
| | time 1 | time 2 | time 3 | time 1 | time 2 | time 3 | time 1 | time 2 | time 3 |
| Heart rate | 68.0 (10.7) | 55.1 (8.7) | 61.1 (9.6) | 68.3 (11.7) | 53.7 (7.9) | 67.9 (11.8) | 68.5 (11.6) | 54.8 (8.4) | 61.3 (10.2) |
| Blood pressure: systolic | 116.2 (10.0) | 112.9 (9.6) | 115.4 (9.0) | 116.7 (8.2) | 114.7 (9.6) | 120.0 (9.9) | 115.9 (9.5) | 114.2 (9.0) | 115.4 (8.6) |
| Blood pressure: diastolic | 62.9 (6.8) | 63.8 (7.5) | 61.8 (6.3) | 62.3 (6.4) | 64.9 (7.4) | 65.4 (7.2) | 61.6 (6.7) | 62.6 (6.4) | 60.9 (5.4) |
| Ear temperature | 35.8 (0.5) | 35.9 (0.4) | 36.1[1] (0.4) | 35.9 (0.5) | 36.0 (0.5) | 36.2 (0.5) | 35.8[1] (0.5) | 36.0 (0.5) | 36.0 (0.5) |
| VAS: medical | 11.7 (3.7) | 11.2 (3.4) | 11.2 (2.6) | 13.4 (4.5) | 12.2 (3.4) | 12.4 (4.1) | 11.5 (3.0) | 11.3 (3.4) | 11.2 (3.2) |
| PANAS: positive affect | 28.9 (6.5) | 27.3 (7.4) | 25.5 (7.8) | 27.8 (7.9) | 27.7 (7.0) | 27.2 (8.3) | 28.1 (7.2) | 27.7 (7.4) | 24.8 (7.7) |
| PANAS: negative affect | 11.5 (2.3) | 11.0 (1.9) | 10.8 (1.5) | 12.0 (2.7) | 11.1 (2.2) | 11.1 (2.1) | 11.8 (2.1) | 11.4 (2.2) | 11.9 (1.9) |
| VAS: alertness | 7.1 (1.5) | 7.2 (1.5) | 6.6 (2.0) | 6.8 (1.6) | 7.2 (1.3) | 7.0 (1.7) | 7.2 (1.4) | 7.1 (1.4) | 6.4 (1.9) |
| VAS: calmness | 7.7 (1.4) | 8.0 (1.4) | 8.0 (1.3) | 7.8 (1.3) | 7.9 (1.4) | 7.5 (1.7) | 7.4 (1.3) | 7.8 (1.6) | 8.0 (1.5) |
| VAS: contentedness | 8.0 (1.1) | 7.9 (1.1) | 7.9 (1.2) | 7.8 (1.4) | 8.0 (1.3) | 8.0 (1.3) | 7.9 (1.3) | 8.1 (1.1) | 7.9 (1.2) |

Mean (SD) scores for heart rate, blood pressure (systolic and diastolic), ear temperature, medical symptoms visual analogue scale and mood measures: Positive and Negative Affect Scale (PANAS); Bond and Lader Visual Analogue Scales (alertness, calmness and contentedness) at each timepoint (baseline, start testing, end testing) for each drug (PBO = placebo; MPH = methylphenidate; SUL = sulpiride). *N* = 46. [1] 1 missing value.

**Inclusion criteria**

Inclusion age range was 18–45 years old and participants had to be native-Dutch speakers, right-handed, had to have normal or corrected-to-normal vision and could not be color-blind. Assessment for inclusion on the first testing day comprised a medical screening, assessing blood pressure (systolic BP: 95-140 mm Hg; dystolic BP: 50-95 mm Hg), heart rate (45-120 bpm) and electrocardiography (QTC-interval M: <450 ms; F: <460 ms; PR-interval: <250 ms), as well as a systematic psychiatric screening interview (M.I.N.I. Plus 5.0.0) assessing psychiatric symptoms, such as major depression, dysthymia, suicidality, (hypo) mania, panic disorder, agoraphobia, social anxiety disorder, obsessive-compulsive disorder, posttraumatic stress disorder, alcohol abuse and dependence, psychoactive substance use disorders, psychotic disorder, anorexia nervosa, bulimia nervosa, generalized anxiety disorder, and attention deficit/hyperactivity disorder. Participants could not have a diagnosis (or history) of relevant psychiatric, neurological, endocrine, or neuroendocrine treatment; frequent autonomic failure; clinically significant hepatic, cardiac, obstructive respiratory, renal, cerebrovascular, cardiovascular, metabolic, ocular or pulmonary diseases/disorders; alcohol or drug dependence; epilepsy; Raynaud's syndrome; one first degree, or two or more second degree family members with history of sudden death of ventricular arrhythmia; history of over the counter medication within the last two months or prescribed medication within the last month prior to the study; regular use of corticosteroids; habitual smoking; diabetes; abnormal hearing or (uncorrected vision); glaucoma; irregular sleep/wake rhythm; possible pregnancy and no appropriate contraception. Participants had to abstain from cannabis throughout the course of the experiment, including 2 weeks before the start of the first pharmacological session, and were required to abstain from alcohol 24 hours and psychotropic medication and recreational drugs 72 hours before each session.

**Detailed description of the behavioral paradigm**

Before drug administration, participants completed a short color sensitivity task (1 minute) and a color wheel working memory task to familiarize participants with the type and level of cognitive control (24 minutes). After drug-intake they completed a cognitive effort-discounting choice task (22 minutes) to quantify the subjective value (and effort costs) of the color wheel working memory task. All tasks were performed on a computer running on Windows 7 and a screen resolution of 1920x1080p. The background color for all tasks was grey (R: 200 G: 200 B: 200). All tasks were programmed in MATLAB version 2016a, using Psychophysics Toolbox Version 3.0.12.

*Color sensitivity task*

The color sensitivity task measured the ability to detect and distinguish the colors in the main task. A color wheel was presented in the middle of the screen and participants had to match the color of a square in the center of the wheel with the corresponding color on the wheel, by clicking on the color wheel. This task consisted of 12 trials and performance on the color sensitivity task was successful if the average deviation of the response from the correct color was below 15 degrees. If participants failed the first time, the color sensitivity task would be assessed again. Participants would be excluded if they failed the color sensitivity task twice (no participants failed the sensitivity test).

The color wheel was created by placing a background-colored circle with a radius of 362p on a circle with a radius of 486p that contained 512 successive colors. By placing the smaller circle over the bigger circle, a colored ring was created. Each color had an angle width of 0.7 degrees and was created with the HSV MATLAB color map. The color of the square was one of the 512 colors from the color wheel. After participants responded, a black line appeared that marked and confirmed the response. The black line consisted of a 0.4-degree arc, placed on the color wheel. Additionally, feedback conveying the deviance (the degrees the response deviated from the correct color) was given if the deviance was below 10 degrees: e.g. 'Good job! You deviated only 4 degrees!' and by a second black arc that marked the correct location of the color. If the deviance was more than 10 degrees, the feedback only consisted of the second black arc. Participants were instructed to answer precisely, but not to take too much time to respond. There was no maximum response time. To test a wide variation of colors from the color wheel, the color wheel was divided into 12 parts, from now on referred to as color pies. From each color pie, one color was chosen at random to be presented in the center of the color wheel. Both the color of the square and the orientation of the color wheel were randomized across participants.

*Color wheel working memory task*

The color wheel task (Figure 1A) is a delayed response task of working memory that distinguishes between distractor resistance and flexible updating, based on a paradigm introduced by Zhang and Luck [19]. Participants had to match a color that was held in working memory to a color wheel on the screen. Each trial was preceded by a centered black dot for 0.5s, which signaled the start of a new trial. On each trial, colored squares were presented in the middle of the screen, with the letter M in the middle, which stood for 'memorize'. During this encoding phase, participants had 0.5s to memorize the colors of the squares, after which there was a delay of 2s. During this delay, a fixation cross was presented in the middle of the screen. Then, during the interference phase, a new set of colored squares appeared on screen, with one of two letters in the center. 'I' stood for 'ignore': participants had to ignore the new squares while still remembering the previous set of squares. 'U' stood for 'update': participants now had to update the colors of the squares, that is, remember the new set of squares and forget the previous squares. The new set of squares remained on screen for 0.5s, followed by a second delay phase. Depending on the task-type, distractor resistance (ignore) or flexible updating (update), this delay lasted either 2s or 4.5s, respectively, to have the same length of time between the relevant stimuli (encoding phase for the ignore type and interference phase for the update type) and the last phase, the probe phase. During this probe phase, a color wheel appeared in the center of the screen, containing frames of colorless squares, one of which was marked. Participants had 4s to indicate the target color of the marked square by clicking on the corresponding color on the color wheel. For example, if the upper right frame was marked, and the condition was ignore, participants had to indicate the color of the upper right square during the encoding phase. When a response was made, a black line appeared on the color wheel at the location where the mouse click was made, remaining on screen until 4s had passed since the appearance of the color wheel. No feedback was given on accuracy. If participants did not respond within 4s, a message was presented for 0.5s in the center of the screen: 'Please respond faster!' Participants were instructed to keep their eyes fixed on the center of the screen during the entire task.

The number and locations of the squares were the same in each phase, but differed over trials, ranging from 1 to 4 squares, allowing us to assess effects of cognitive load (from now on referred to as set-size). All combinations of set-size (ranging from 1-4) and task-type (ignore or update) were repeated 16 times, which resulted in 128 trials divided over two blocks. The task was preceded by 16 practice trials. On these practice trials, feedback conveying the deviance (the degrees the response deviated from the correct color, with a maximum of 180 degrees) was given if the deviance was below 10 degrees: e.g. 'Good job! You deviated only 4 degrees!' and by a second black line that marked the correct location of the color. If the deviance was more than 10 degrees, the feedback only consisted of the second black line. Feedback duration was 0.7s. To control for possible effects of different colors, the colors that were presented during the encoding phase of ignore trials were the same as the colors that were used during the interference phase of update trials. Additionally, the target colors were the same for both conditions. To control for possible location effects, the locations of the squares were allocated equally across trials and the location of the marked frame, and thus the target, was balanced across conditions. Additionally, the orientation of the color wheel was randomized across trials. Trial order was the same for each participant.

*Choice task*

To quantify participant's preference for the task versus rest, participants completed a choice task where they repeatedly chose between a cognitively effortful (redo) option for more money and a leisure (no-redo) option for less money (Figure 1B). A redo choice implied that they preferred to complete another block of the color wheel task after completing the choice task. By choosing the no-redo option they indicated that they preferred to be free to do what they wanted, such as using their phone or the computer, for an equal length of time as another round of the color wheel task, while staying in the testing room. Participants were informed that their monetary bonus and the difficulty of the working memory task would depend on their choices during the choice task, because one of their choices would be randomly selected. Due to time constraints and to avoid transfer effects of experiencing the color wheel task under drug to future sessions, both the monetary bonus and the redo of the color wheel task were hypothetical, and participants were instructed accordingly. The rationale for this instruction was that it pre-empted gradual learning that the redo phase was hypothetical, invalidating comparison between the three sessions. Despite its hypothetical nature, the task manipulation and sensitivity to cognitive effort was validated by evidence for strong monotonic, set size-dependent discounting (see Results). The hypothetical compensation for the redo option was fixed at €2.00. The compensation for the no-redo option varied from a minimum of €0.10, and then from €0.20 to €2.20, with intervals of €0.20. The redo option was further specified by a task-type (ignore or update) and a set-size (1-4), which meant that most of the redo block would consist of trials of that task-type and set-size. The remainder of the trials would be divided among all task-type and set-size combinations. The task was divided into three blocks, with a break in between the blocks. Each block featured each unique combination of task-type, set-size and monetary compensation for the no redo option (€0.10-€2.20) in random order, which resulted in 96 trials per block and 288 trials in total. On each trial participants had 4s to respond. They were told that to receive the bonus, their performance during the redo block would have to be similar to their performance during the earlier blocks and that this meant that they had to put effort into doing the redo block, but not that they always had to be correct. This was conveyed to minimize differences in preference for a condition due to earlier performance differences.

### SUPPLEMENTAL RESULTS

#### Voxel-wise PET analyses

We conducted voxel-wise analyses in addition to our primary region-of-interest analyses to assess the physiological plausibility of the link between drug effects on choice behavior (proportion redo choices and choice latency) and striatal dopamine synthesis capacity ($K_i$ influx constant). To this end, the individual $K_i$ maps were spatially normalized to MNI space and smoothed using an 8mm FWHM kernel. We restricted our search, following prior procedures [20], to one region of interest comprising all voxels that exhibited a $K_i$ value of 3 standard deviations above the global mean. This region of interest included the striatum and midbrain (8993 voxels; Figure S1). Statistical significance was defined as family-wise error (FWE) corrected $p<0.05$ at peak coordinate, after small volume correction for all voxels within the region of interest. Note that the group-based [$^{18}$F]DOPA mask was calculated based on the larger study group excluding drop-outs (*N* = 94).

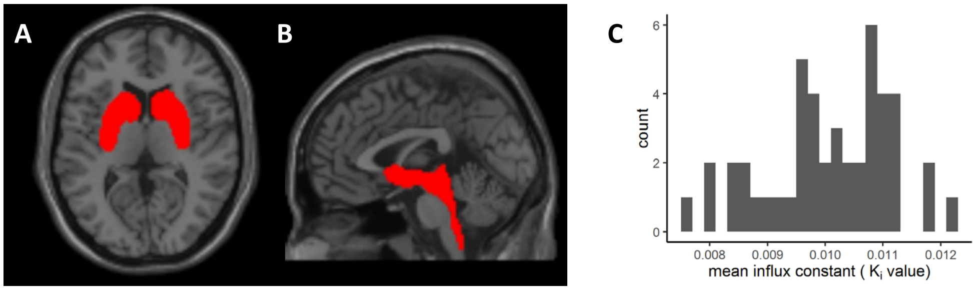


**Figure S1**. **A** – Axial and **B** – sagittal view of the group-based small volume mask including all voxels with a $K_i$ value of > 3 SD from the global mean. **C** – Histogram of mean $K_i$ value within the group-based mask. Count represents number of participants. N = 46. The mean $K_i$ values varied between 0.00768 and 0.01228. These values are comparable with previous reports [21].

High-dopamine participants exhibited greater methylphenidate-related increases in cognitive motivation

In accordance with our region-of-interest based results, voxel-wise analyses revealed a positive correlation between baseline dopamine synthesis capacity and the methylphenidate effect on overall proportion redo choices (Figure S2B; Table S6). Participants with higher dopamine synthesis capacity exhibited greater methylphenidate-induced increases in proportion redo choices than participants with lower dopamine. The correlation with midbrain dopamine synthesis capacity survived *FWE*-correction. Although not surviving *FWE*-correction, positive correlations with the methylphenidate effect were also found in the nucleus accumbens, putamen, and caudate nucleus. There was no association between the effect of sulpiride on proportion redo choices and baseline dopamine synthesis capacity (Figure S2C) and no voxels correlated significantly with the difference in proportion redo choices between methylphenidate and sulpiride (Figure S2D; Table S6). The correlation between baseline dopamine synthesis capacity and proportion redo choices under placebo was not significant (Figure S2A; Table S6).

High-dopamine participants exhibited greater methylphenidate-related slowing of choice latency

The effect of methylphenidate on choice latency correlated positively with baseline dopamine synthesis capacity in the midbrain, left caudate nucleus and right putamen (Figure S2F; Table S6), indicating that methylphenidate slowed people with higher baseline dopamine synthesis capacity and invigorated people with lower baseline dopamine synthesis capacity. No significant correlation between baseline dopamine synthesis capacity and the effect of sulpiride was observed (Figure S2G; Table S6) and no voxels correlated significantly with the difference in choice latency on the methylphenidate and the sulpiride session (Figure S2H; Table S6). Dopamine synthesis capacity correlated negatively with choice latency under placebo in the left caudate nucleus and midbrain (Figure S2A; Table S6), indicating that higher baseline dopamine synthesis capacity was associated with faster responding.

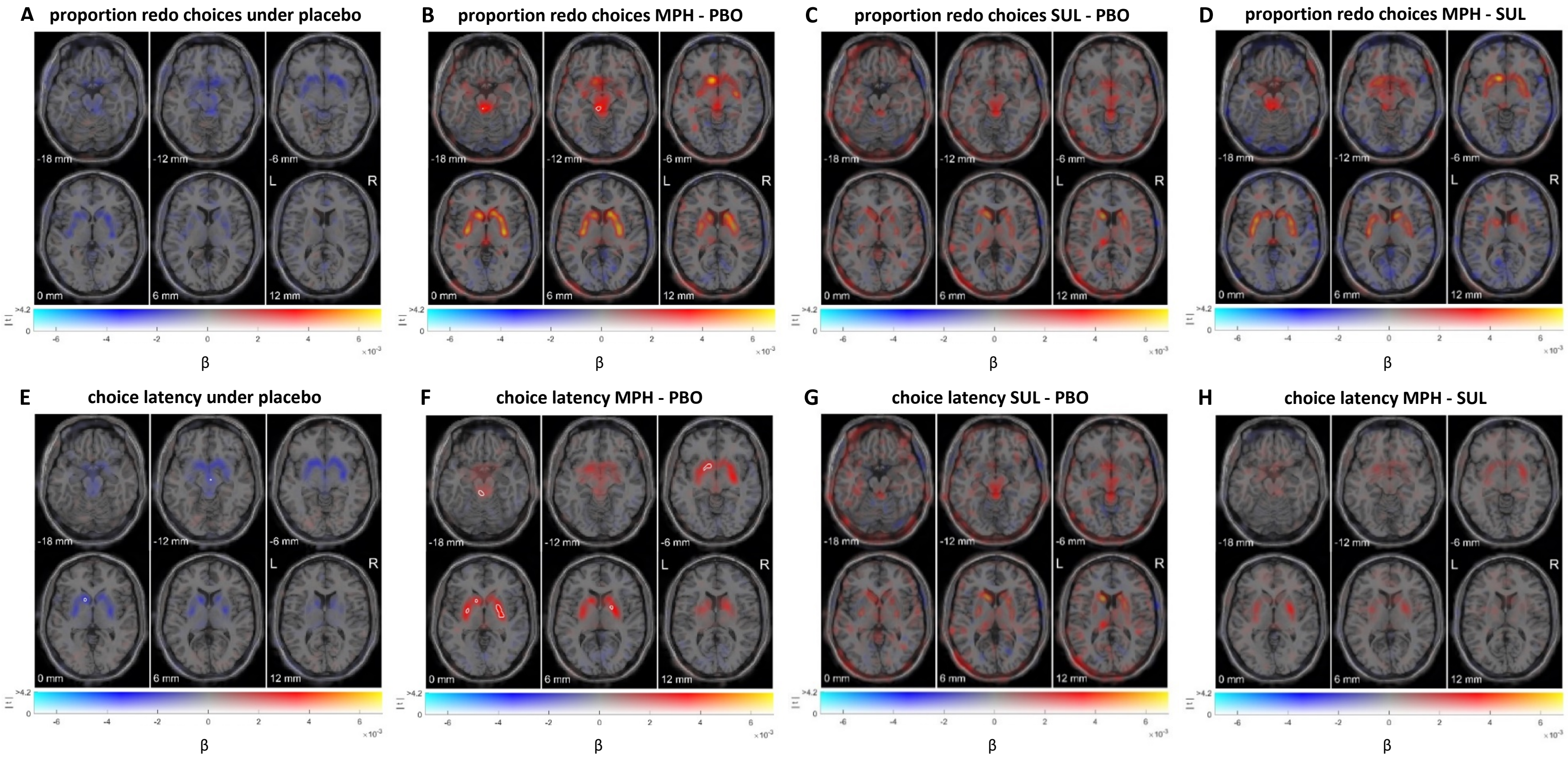


**Figure S2.** Voxels showing a positive (red) or negative (blue) regression coefficient on **(A)** the proportion redo choices under placebo (PBO), **(B)** the effect of methylphenidate (MPH) versus placebo on proportion redo choices, **(C)** the effect of sulpiride (SUL) versus placebo on proportion redo choices, **(D)** the effect of methylphenidate versus sulpiride on proportion redo choices, **(E)** choice latency under placebo, **(F)** the effect of methylphenidate versus placebo on choice latency, **(G)** the effect of sulpiride versus placebo on choice latency, **(H)** the effect of methylphenidate versus sulpiride on choice latency. The brain maps are dual-coded and simultaneously display the contrast estimate (*x* axis) and *t* values for the drug contrast (*y* axis). The hue indicates the size of the contrast estimate, and the opacity indicates the height of the *t* value. Significant clusters (peak-level corrected, FWE, $p<0.05$) are encircled in white. The *z* coordinates correspond to the standard MNI brain. Neuroimaging data are plotted using a procedure introduced by Allen et al. [22] and implemented by Zandbelt [23].

**Table S6. Brain regions exhibiting correlations between dopamine synthesis capacity, and choice and working memory performance**

| | | | | | | coordinates | | |
|---|---|---|---|---|---|---|---|---|
| *Contrast* | *anatomical region* | *hemisphere* | *direction* | *t* | *corrected p* | *x* | *y* | *z* |
| Proportion redo choices PBO | midbrain | right | negative | 3.96 | 0.099 | 6 | -36 | -14 |
| **MPH effect on proportion redo choices** | **midbrain** | **left** | **positive** | **4.73** | **0.013** | **-4** | **-32** | **-14** |
| | Pons | left | positive | 3.86 | 0.124 | -4 | -38 | -34 |
| | nucleus accumbens | left | positive | 3.68 | 0.185 | -6 | 14 | -6 |
| | caudate nucleus | right | positive | 3.47 | 0.283 | 20 | 6 | 14 |
| | caudate nucleus | right | positive | 3.32 | 0.373 | 16 | -6 | 20 |
| MPH vs SUL effect on proportion redo choices | midbrain | left | positive | 3.55 | 0.254 | -8 | -30 | -16 |
| **MPH effect on indifference point** | **midbrain** | **left** | **positive** | **4.65** | **0.017** | **-4** | **-30** | **-16** |
| **Choice latency PBO** | **caudate nucleus** | **left** | **negative** | **4.44** | **0.029** | **-10** | **12** | **0** |
| | **midbrain** | **right** | **negative** | **4.27** | **0.045** | **4** | **-14** | **-12** |
| | midbrain | right | negative | 4.11 | 0.068 | 4 | -36 | -14 |
| | putamen | right | negative | 3.46 | 0.287 | 32 | -6 | -4 |
| | putamen | right | negative | 3.45 | 0.290 | 20 | 18 | -8 |
| | amygdala | right | negative | 3.34 | 0.357 | 28 | -2 | -14 |
| **MPH effect on choice latency** | **putamen** | **right** | **positive** | **5.03** | **0.006** | **28** | **-8** | **0** |
| | **caudate nucleus** | **left** | **positive** | **4.94** | **0.007** | **-10** | **10** | **-4** |
| | **midbrain** | **left** | **positive** | **4.77** | **0.012** | **-4** | **-34** | **-18** |
| | midbrain | left | positive | 3.51 | 0.258 | -6 | -8 | -10 |
| | medulla oblongata | right | positive | 3.50 | 0.266 | 2 | -42 | -46 |
| | caudate nucleus | left | positive | 3.48 | 0.274 | -18 | -2 | 14 |
| | nucleus accumbens | right | positive | 3.39 | 0.326 | 4 | 12 | -12 |
| SUL effect on choice latency | caudate nucleus | left | positive | 3.68 | 0.183 | -10 | 8 | -4 |
| | caudate nucleus | right | positive | 3.50 | 0.266 | 18 | 4 | 12 |
| MPH vs SUL effect on choice latency | putamen | right | positive | 3.59 | 0.236 | 28 | 0 | 0 |
| | Pons | right | positive | 3.41 | 0.334 | 6 | -38 | -24 |
| | midbrain | right | positive | 3.36 | 0.364 | 8 | -32 | -18 |
| | amygdala | left | positive | 3.34 | 0.380 | -18 | -6 | -16 |
| MPH effect on deviance (working memory task) | midbrain | right | positive | 3.33 | 0.248 | 4 | -38 | -20 |
| | caudate nucleus | left | positive | 3.25 | 0.299 | -12 | 6 | 2 |
| | midbrain | right | positive | 3.18 | 0.350 | 2 | -36 | -14 |
| | caudate nucleus | left | positive | 3.17 | 0.355 | -18 | 0 | 16 |

Spatial coordinates of local maxima for regions showing a correlation with behavioral measures. Coordinates correspond to the standard MNI brain. Statistical interference was based on a peak-level correction of $p < 0.05$ (FWE) using small volume correction (a volume comprising all voxels with a $K_i$ value of > 3 SD from the global mean). Caution is warranted: shown here, for full reporting, are all clusters with an uncorrected *p*-value < 0.001. Significant clusters are in bold.
PBO: placebo; MPH: methylphenidate; SUL: sulpiride. N = 46.

**Effects on proportion redo choices are reproduced when analysing indifferent points**

A priori we had planned to calculate indifference points (IPs): The offer for the no-redo option at which the participant was indifferent to either choosing no-redo or redoing the task for €2.00 (Figure S3A). However, because many participants demonstrated a very low willingness to redo the task, sometimes resulting in inestimable IPs, we instead decided to base our primary analyses on the proportion redo choices. A high IP means a strong preference for the cognitively more demanding redo option and high motivation for task engagement. The IP was calculated using binomial logistic regression analysis. Where IP was lower or higher than the minimum (€0.10) or maximum (€2.20) offer for the no-redo option, respectively, the IP was set to either 0.10 (in case of an IP lower than the minimum) or 2.20 (in case of an IP higher than the maximum), to avoid loss of valuable data. Table S7 shows the repeated-measures ANOVA results with the within-participants factors drug (methylphenidate, sulpiride, placebo), task-type (ignore, update) and set-size and the covariate dopamine synthesis capacity (separate analyses for the three ROIs – nucleus accumbens, putamen and caudate nucleus).

At baseline, participants exhibited a strong preference for not repeating the task, as evidenced by the overall IP being significantly smaller than €2.00, the fixed offer for the redo option, on the placebo session (0.79, SD = 0.55; $t_{(45)} = -14.9$, $p < 0.001$).

In accordance with proportion redo choices, IP decreased with set-size ($F_{(1.37,60.29)} = 103.3$, $p < 0.001$). There was no effect of task-type ($F_{(1,44)} = 1.6$, $p = 0.216$) and no interaction between task-type and set-size ($F_{(2.28,100.25)} = 0.5$, $p = 0.622$). We found a significant effect of drug on IP (main effect over the 3 drug conditions: $F_{(2,88)} = 6.4$, $p = 0.003$), which was driven by a positive effect of methylphenidate versus placebo ( $F_{(1,44)} = 9.2$, $p = 0.004$; Figure S3B) and a positive effect of methylphenidate versus sulpiride ( $F_{(1,44)} = 8.6$, $p = 0.005$). There was no difference between sulpiride and placebo ($F_{(1,44)} = 0.1$, $p = 0.780$; Figure S3B).

The interaction between drug effect on IP and dopamine synthesis capacity in the nucleus accumbens and putamen tended towards significance (Table S7). Again, this was driven by a difference between methylphenidate and placebo, where participants with higher dopamine synthesis capacity exhibited greater methylphenidate-induced increases in IP than participants with lower dopamine synthesis capacity. A similar trend was present for the difference between methylphenidate and sulpiride, with participants with higher dopamine synthesis capacity exhibiting greater methylphenidate-induced than sulpiride-induced increases in IP than participants with lower dopamine synthesis capacity. The difference between sulpiride and placebo was not significant. A negative association between baseline dopamine synthesis capacity and IP under placebo was, although trending in the nucleus accumbens, not significant (Table S7).

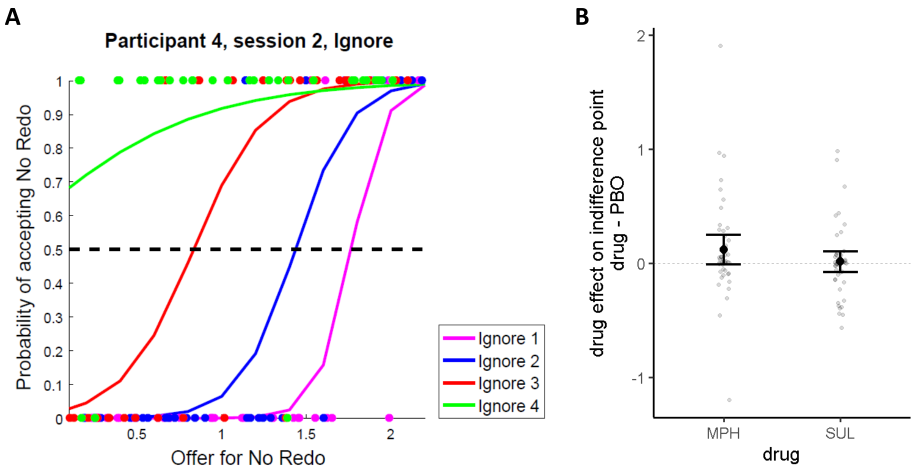


**Figure S3**. **Indifference point. A –** Probability of choosing the no-redo option as a function of the monetary offer for the no-redo option on Ignore trials during session 2 for a representative participant. Fitted probability functions for each set-size are plotted using logistic regression. The indifference point (IP) – where the probability line hits the dashed line – is the offer for the no-redo option at which there is a 50% chance that the participant chose no-redo for that amount or redo for €2. **B** – Drug effect on indifference point (methylphenidate or sulpiride minus placebo). Error bars represent 95% confidence interval around the mean. MPH: methylphenidate; SUL: sulpiride; PBO: placebo.

**Table S7. Repeated-measures ANOVAs on indifference points.** Separate analysis for each ROI – nucleus accumbens, putamen and caudate nucleus, including drug, set-size and task-type as within-subjects variables and baseline dopamine synthesis capacity (measured as the mean-centered average [$^{18}$F]DOPA uptake, $K_i$) as covariate. Statistics for the interaction between baseline dopamine synthesis and drug are shown, as well as the main effect of dopamine synthesis capacity on the placebo session. MPH = methylphenidate; SUL = sulpiride; PBO = placebo. *p*-values below a Bonferroni-corrected alpha-value of 0.017 were considered significant.

| | **Nucleus accumbens** | | **Putamen** | | **Caudate nucleus** | |
|---|---|---|---|---|---|---|
| | $F_{(2,88)}$ | *p* | $F_{(2,88)}$ | *p* | $F_{(2,88)}$ | *p* |
| MPH, SUL, PBO | 3.4 | 0.036 | 3.2 | 0.044 | 2.2 | 0.112 |
| *post-hoc:* | $F_{(1,44)}$ | *p* | $F_{(1,44)}$ | *p* | $F_{(1,44)}$ | *p* |
| MPH, PBO | 5.6 | 0.022 | 5.4 | 0.025 | 3.9 | 0.054 |
| SUL, PBO | 0.5 | 0.502 | 0.6 | 0.451 | 1.7 | 0.198 |
| MPH, SUL | 3.5 | 0.067 | 3.0 | 0.088 | 0.8 | 0.366 |
| | $F_{(1,44)}$ | *p* | $F_{(1,44)}$ | *p* | $F_{(1,44)}$ | *p* |
| PBO | 3.8 | 0.058 | 1.5 | 0.221 | 0.3 | 0.619 |

**Methylphenidate-induced effect does not reflect choice randomness**

It is possible that for high-dopamine individuals, methylphenidate increased choice randomness rather than cognitive motivation. Increased choice randomness would bring the proportion redo choices closer towards 0.5, which would, at the same time, increase the proportion redo choices. We therefore explored the choice slopes (see also Figure S3A). If participants were more random in their choices, their slope should be shallower. One person had an overall slope that was higher than three standard deviations above the global mean and was therefore excluded. Moreover, since the slope could not be estimated in case the IP was below the minimum of €0.10, we restricted the within-subjects variables to the factor drug (with conditions methylphenidate and placebo) to maximize the number of participants that could be included. This resulted in N = 33. A rmANOVA revealed no effect of drug ($F_{(1,30)} = 2.2$, $p = 0.150$) nor an interaction between dopamine synthesis capacity and drug (nucleus accumbens: $F_{(1,30)} = 0.1$, $p = 0.710$; putamen: $F_{(1,30)} = 0.02$, $p = 0.880$; caudate nucleus: $F_{(1,30)} = 0.0$, $p = 0.997$). We thus concluded that the baseline dopamine-dependent effect of methylphenidate on motivation did not reflect an effect on choice randomness.

**Baseline dopamine-dependent effects cannot be explained by working-memory performance**

Although drugs were only administered after the working-memory task, we wanted to verify that effects on performance could not explain the baseline dopamine-dependent effects of methylphenidate. We therefore ran the rmANOVAs again for deviance and RT, including the within-subjects variables task-type, set-size and drug and the covariate dopamine synthesis capacity. There was no interaction between dopamine synthesis capacity and drug on deviance (nucleus accumbens: $F_{(2,88)} = 1.9$, $p = 0.152$; putamen: $F_{(2,88)} = 2.8$, $p = 0.069$; caudate nucleus: $F_{(2,88)} = 1.4$, $p = 0.258$) or RT (nucleus accumbens: $F_{(2,88)} = 0.0$, $p = 0.991$; putamen: $F_{(2,88)} = 0.4$, $p = 0.643$; caudate nucleus: $F_{(2,88)} = 0.0$ $p = 0.991$). The main effect of drug was also not significant (deviance: $F_{(2,88)} = 1.2$, $p = 0.307$; RT: $F_{(2,88)} = 0.8$, $p = 0.453$). No other interactions including drug or dopamine synthesis capacity were significant.

We also assessed a potential association between methylphenidate-related changes in performance and proportion redo choices or choice latency with a Pearson's correlation. As expected, the effect of methylphenidate on performance did not correlate with the effect of methylphenidate on proportion redo choices (deviance: $r = 0.10$, $p = 0.493$; RT: $r = 0.16$, $p = 0.276$) or choice latency (deviance: $r = 0.16$, $p = 0.297$; RT: $r = 0.14$, $p = 0.347$). It is therefore unlikely for methylphenidate-effects on choice behavior to be explained by performance.

**Baseline dopamine-dependent effects cannot be explained by mood and medical symptoms**

On the pharmacological sessions (day 2-4), blood pressure, heart rate and ear temperature were monitored three times on each session: before the start of the task battery, 20 minutes before the start of the discounting task and after the task battery. At the same timepoints, mood measures (Positive and Negative Affect Scale [24]; the Bond and Lader Visual Analogue Scales (calmness, contentedness, alertness; [25])) and medical symptoms (medical visual analogue scale) were assessed. Table S5 gives an overview of those measures for the participants included in the analyses.

To assess whether methylphenidate-induced effects could be accounted for by nonspecific effects of methylphenidate on mood and medical symptoms, we performed a repeated measures MANOVA in SPSS (IBM SPSS statistics version 23.0) using Pillai's trace with the within-subject factors Time (3: before the start of the task battery, 20 minutes after intake of the second capsule and after the task battery) and Drug (3: placebo, methylphenidate and sulpiride) and the six measures as dependent variables (positive affect, negative affect, calmness, alertness, contentedness, medical symptoms). Significant initial MANOVA results ($p < 0.05$) were followed up with univariate tests of the interaction effect (drug x time) for each of the six dependent variables, with Bonferroni-corrected alpha = 0.05/6 ≈ 0.008.

Drug did not significantly affect the mood and medical measures over time (drug x time: $V = 0.68$, $F_{(24,22)} = 2.0$, $p = 0.057$). We conducted the MANOVA again, but now with only the first two timepoints, given the proximity of the second timepoint to the choice task (20 minutes before the start of the task). This revealed a significant drug effect over time (drug x time: $V = 0.48$, $F_{(12,34)} = 2.6$, $p = 0.015$). However, none of the univariate interactions between drug and time for the six dependent variables was significant (all $p > 0.061$). The effect was driven by a difference between the methylphenidate- and sulpiride-effects over time (drug x time; 2 timepoints: $V = 0.34$, $F_{(6,40)} = 3.4$, $p = 0.008$), with a trend for methylphenidate but not sulpiride increasing alertness over time ($F_{(1,45)} = 4.8$, $p = 0.034$) and decreasing medical symptoms over time ($F_{(1,45)}$ = 4.4, $p$ = 0.041). The drug by time interactions for the other dependent variables were all above $p$ = 0.143). A MANOVA including only sulpiride and placebo showed a difference of the drug effects over time (drug x time; 2 timepoints: $V$ = 0.28, $F_{(6,40)}$ = 2.6, $p$ = 0.035). However, none of the univariate interactions between drug and time for the six dependent variables was significant (all $p$ > 0.084). Importantly, the methylphenidate-effect over time did not differ from that of placebo (drug x time; 2 timepoints: $V$ = 0.20, $F_{(6,40)}$ = 2.6, $p$ = 0.160).

A MANOVA on only timepoint 2 revealed no significant effect of drug ($V$ = 0.38, $F_{(12,34)}$ = 1.7, $p$ = 0.107). Next, we correlated methylphenidate-induced changes on the six mood and medical measures at timepoint 2 with methylphenidate-induced changes in proportion redo choices. This revealed a significant positive correlation for positive affect ($r$ = 0.34, $p$ = 0.022) and a trending positive correlation for alertness ($r$ = 0.28, $p$ = 0.60). The other correlations were all above $p$ = 0.283. We therefore ran additional repeated-measures ANOVAs on proportion redo choices with drug (methylphenidate and placebo), task-type (ignore and update) and set-size as within-subjects variables and dopamine synthesis capacity as a covariate, while also including the effect of methylphenidate on positive affect and alertness, respectively, as a covariate. Methylphenidate still significantly affected proportion redo choices when including positive affect, both as a main effect ($F_{(1,43)}$ = 9.0, $p$ = 0.005) and in interaction with dopamine synthesis capacity (nucleus accumbens: $F_{(1,43)}$ = 6.7, $p$ = 0.013; putamen: $F_{(1,43)}$ = 7.7, $p$ = 0.008; caudate nucleus: $F_{(1,43)}$ = 5.4, $p$ = 0.024). Similarly, methylphenidate still significantly affected proportion redo choices when including alertness, both as a main effect ($F_{(1,43)}$ = 9.2, $p$ = 0.004) and in interaction with dopamine synthesis capacity (nucleus accumbens: $F_{(1,43)}$ = 11.3, $p$ = 0.002; putamen: $F_{(1,43)}$ = 11.0, $p$ = 0.002; caudate nucleus: $F_{(1,43)}$ = 7.9, $p$ = 0.008). It is therefore unlikely that drug-induced changes on mood or medical measures account for effects on proportion redo choices.

**Controlling for session order**

High-dopamine individuals received methylphenidate on earlier sessions than low-dopamine individuals

A first ANOVA confirmed that session-day-number (1, 2 or 3) did not differ between the three drugs ($F_{(2, 144)}$ = 0.9, $p$ = 0.392). The order of the drug session could further potentially have confounded our baseline-dependency effects if people with lower dopamine synthesis capacity accidentally experienced a different order than people with higher synthesis capacity. Unexpectedly, a Spearman's rank correlation indeed revealed that dopamine synthesis capacity was negatively correlated with the session on which participants received methylphenidate (nucleus accumbens: rho = -0.29, $p$ = 0.049; putamen: rho = -0.40, $p$ = 0.005; caudate nucleus: rho = -0.24, $p$ = 0.116; Table S8). Participants with higher dopamine synthesis capacity more often received methylphenidate on the earlier sessions. There were no significant correlations between dopamine synthesis capacity and the session on which participants received sulpiride or placebo (Table S8). Given these findings, we reanalyzed our significant methylphenidate effects and methylphenidate by baseline dopamine synthesis capacity interactions while controlling for the session number for methylphenidate by adding methylphenidate's session number as an ordinal between-subjects factor to the rmANOVA.

Baseline dopamine synthesis capacity-dependent effects are smaller, but still present

After including methylphenidate's session number as a between-subjects factor, methylphenidate, relative to placebo, indeed still significantly increased proportion redo choices ($F_{(1,42)}$ = 6.9, $p$ = 0.012). The interaction between dopamine synthesis capacity and the effect of methylphenidate on proportion redo choices was, although smaller, still present (nucleus accumbens: $F_{(1,42)}$ = 4.6, $p$ = 0.038; putamen: $F_{(1,42)}$ = 2.4, $p$ = 0.127; caudate nucleus: $F_{(1,42)}$ = 4.0, $p$ = 0.051). Similarly, the interaction between dopamine synthesis capacity and the effect of methylphenidate on choice latency was also still present, but smaller (nucleus accumbens: $F_{(1,42)}$ = 5.3, $p$ = 0.026; putamen: $F_{(1,42)}$ = 3.4, $p$ = 0.071; caudate nucleus: $F_{(1,42)}$ = 1.3, $p$ = 0.252).

<u>No correlation between dopamine synthesis capacity and time after methylphenidate administration</u>

Previous research suggested that increased striatal dopamine synthesis capacity can be detected by an [$^{18}$F]DOPA scan in conjunction with the inlet–outlet model even 2 weeks after a methylphenidate treatment [26]. We therefore assessed whether the order effects could have arisen due to too short intervals between the drug sessions and the PET scan. The interval between the methylphenidate session and the PET scan ranged between 7 and 106 days (mean = 43.0, SD = 26.8). However, Pearson's correlations between dopamine synthesis capacity and days between the methylphenidate session and the PET scan were not significant (nucleus accumbens: $r$ = 0.16, $p$ = 0.; putamen: $r$ = 0.18, $p$ = 0.245; caudate nucleus: $r$ = 0.04, $p$ = 0.786), thus minimizing the likelihood that our index of dopamine synthesis capacity was affected by drug administration.

**Table S8.** Spearman's rank correlations between dopamine synthesis capacity and session number on which participants received methylphenidate, sulpiride and placebo.

| | **Nucleus accumbens** | | **Putamen** | | **Caudate nucleus** | |
|---|---|---|---|---|---|---|
| | *rho* | *p* | *rho* | *p* | *rho* | *p* |
| Methylphenidate | -0.29 | 0.049 | -0.40 | 0.005 | -0.24 | 0.116 |
| Sulpiride | 0.11 | 0.473 | 0.24 | 0.107 | 0.11 | 0.453 |
| Placebo | 0.21 | 0.153 | 0.21 | 0.167 | 0.15 | 0.310 |

**Drug effects on physiological measures**

Primary analyses revealed no significant effects of sulpiride. To assess whether this could be explained by a general lack of sulpiride effect, we performed a repeated measures MANOVA on the physiological measures heart rate, systolic blood pressure, diastolic blood pressure and ear temperature. The within-subject factors were Time (3: before the start of the task battery, 20 before the start of the discounting task and after the task battery) and Drug (3: placebo, methylphenidate and sulpiride). Significant effects were followed up with univariate tests of the interaction effect (drug x time) for each of the four dependent variables, with Bonferroni-corrected alpha = 0.05/4 ≈ 0.013.

Drug significantly affected heart rate and blood pressure over time (drug x time: $V = 0.73$, $F_{(16,29)} = 4.9$, $p < 0.001$; heart rate: $F_{(4,176)} = 14.5$, $p < 0.001$; systolic blood pressure: $F_{(4,176)} = 6.5$, $p < 0.001$; diastolic blood pressure: $F_{(4,176)} = 6.4$, $p < 0.001$), with a trend for ear temperature ($F_{(3.15,138.46)} = 2.9$, $p = 0.035$). The effect was driven by a difference between methylphenidate and placebo ($V = 0.65$, $F_{(8,37)} = 8.6$, $p < 0.001$). Over time, methylphenidate increased heart rate ($F_{(2,88)} = 20.4$, $p < 0.001$), systolic blood pressure ($F_{(2,88)} = 7.4$, $p = 0.001$), diastolic blood pressure ($F_{(1.74,76.46)} = 9.2$, $p < 0.001$) and, sub-threshold, ear temperature ($F_{(1.76,77.20)} = 3.4$, $p = 0.046$) compared with placebo. There were no significant effects of sulpiride over time ($V = 0.14$, $F_{(8,37)} = 0.7$, $p = 0.654$). We conducted the MANOVA again, but now with only the first two timepoints, given the proximity of the second timepoint to the choice task (20 minutes before the start of the task). This revealed no significant drug effect over time (drug x time: $V = 0.18$, $F_{(8,37)} = 1.0$, $p = 0.426$). A MANOVA on only timepoint 2 revealed a significant effect of drug ($V = 0.44$, $F_{(8,38)} = 3.7$, $p = 0.003$). However, only the univariate analysis on diastolic blood pressure revealed a non-significant trend ($F_{(2,90)} = 3.9$, $p = 0.023$). The interactions for the other dependent variables were all above $p = 0.145$. The trend-level interaction on diastolic blood pressure stemmed from a difference between methylphenidate and sulpiride ($F_{(1,45)} = 7.1$, $p = 0.145$), where diastolic blood pressure was higher on the methylphenidate session. Thus, although there were clear physiological effects of methylphenidate over time, these only emerged when including the third timepoint. There were no clear physiological effects of sulpiride.

### SUPPLEMENTAL DISCUSSION

#### Cognitive stability versus cognitive flexibility

In addition to investigating choices between cognitive control and rest, we assessed choices between different types of cognitive control. Specifically, we were interested to explore catecholaminergic drug effects on choices between a working memory “update” task requiring cognitive flexibility and a working memory “ignore” task requiring cognitive stability [27–29]. Our data showed no strong evidence for methylphenidate or sulpiride shifting the balance between the motivation for stable versus flexible control. This might be not surprising, because, contrary to earlier studies, in which participants preferred a similar flexible update task over a stable ignore task [15, 30], there was no difference between the value of the tasks under placebo. It is potentially relevant that here, in contrast to that previous work, participants did not choose between the “update” or “ignore” task, but rather between one of those tasks and a rest option. For future work, to investigate the question whether dopamine alters preference for one task versus another, we would require direct choices between the two types of task, while also considering the manipulation across time of the frequency of one task-type over another.

22. Allen EA, Erhardt EB, Calhoun VD. Data visualization in the neurosciences: overcoming the curse

of dimensionality. Neuron. 2012;74:603–608.
